# Design to Data for mutants of β-glucosidase B from *Paenibacillus polymyxa*: M319C, T431I, and K337D

**DOI:** 10.1101/839027

**Authors:** Peishan Huang, Stephanie C. Contreras, Eliana Bloomfield, Kristine Schmitz, Augustine Arredondo, Justin B. Siegel

## Abstract

The use of computational tools has become an increasingly popular tool for engineering protein function. While there are numerous examples of computational tools enabling the design of novel protein functions, there remains room for improvement in both prediction accuracy and success. To improve algorithms for functional and stability predictions, we have initiated the development of a data set designed to be used for training new computational algorithms for enzyme design. To date our dataset is composed of over 129 mutants with associated expression levels, kinetic data, and thermal stability for the enzyme β-glucosidase B (BglB) from *Paenibacillus polymyxa*. In this study, we introduced three new variants (M319C, T431I, and K337D) to our existing dataset with the goal of cultivating a larger dataset to train new design algorithms and more broadly explore structure-function relationships in BglB.

## INTRODUCTION

Computational tools to engineer protein catalytic efficiency, stability, or substrate specificity are becoming crucial for the development of industrial and therapeutic proteins. Despite recent success in using rational design for thermal stability,^1–4^ or creating *de novo* enzymes^5–8^ not found in nature, current algorithms are inconsistent in their performance on different biological systems and often have a lower success than desirable.^9^

One potential reason for inconsistent and low prediction accuracies is that the current algorithms are trained primarily on protein structure as opposed to function. This is particularly challenging for enzymes, many of which are believed to adopt multiple conformations to accommodate the reaction from of a substrate product.^11^ Furthermore, for enzymes, there are no large datasets that quantitatively measure structure-function relationships on potentially modellable physical properties (e.g. T_M_, *k*_cat_, K_M_, and *k*_cat_/K_M_). Therefore, we have initiated an effort to systematically characterize a family 1 glycoside hydrolase (GH1) from *Paenibacillus polymyxa*, Beta-glucosidase B (BglB, Uniprot ID: P22505). Here we present three new variants (M319C, T431I, and K337D) to explore the mutational effects surface residues have on kinetic activity and thermal stability. To date, our dataset consists of over 129 mutants that have been characterized for T_50_, *k*_cat_, K_M_, and *k*_cat_/K_M12_ and we have added these three new variants to continue the expansion of our dataset for training an algorithm to have better predictive functional properties and thermal stability. Currently, the majority of the variants are found near the active-site of the BglB, to explore the mutational effects of surface residues on kinetic activity and thermal stability, M319C, T431I, and K337D (Fig 1) were chosen for testing.

**Figure 1.**
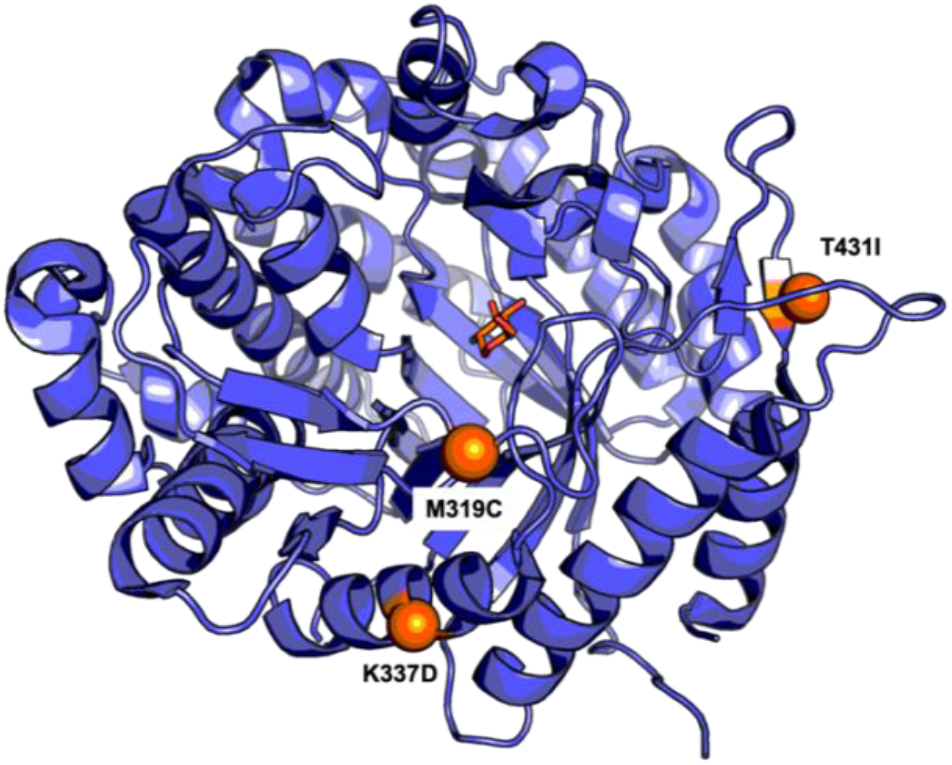
Rendering of BglB from *Paenibacillus polymyxa*. The three mutational sites that were made in study are highlighted in orange spheres on the cartoon illustration of BglB (PDB ID: 2JIE), rendered using PyMOL.^10^

## METHODS

### Design and building of new variants

Using a previously described method,^9^ three single-point mutations, M319C, T431I, and K337D were selected and modeled using FoldIt.^13^ The three surface mutational changes were scored by the Rosetta energy function and are given a total system energy score (TSE).^14^ No mutants were selected if the changes between the wild type and variant score is predicted to be greater than +5 to enrich for proteins likelihood to fold and function.

### Generating mutant plasmids

The mutant plasmids were generated by Kunkel mutagenesis^15^ as previously described.^12^ The plasmids were electroporated into Dh5*α* competent *Escherichia coli* (E.*coli*) cells and plated onto Luria-Bertani (LB) agar plate with 50 *μg*/mL kanamycin for overnight growth. Single colonies were picked and sequence to identify the desired mutants.

### Protein production and purification

The sequence-verified BglB mutant DNA were transformed, grown, and expressed using the previously described method.^12^ Briefly, following autoinduction the cells were lysed and proteins purified using immobilized metal ion affinity chromatography. The protein purity was analyzed using 4-12% SDS-PAGE (Life Technologies) and all samples used in the study were > 80 % pure. The protein concentrations were determined by measuring absorbance at 280 nm and calculating the expected molarity based on the predicted extinction coefficient from ProtParam.^16^

### Michaelis-Menten kinetics and temperature melting assays (T_M_)

The kinetic characterization was done in the same manner as previous study.^9^ Briefly, the activity was obtained by monitoring the production of 4-nitrophenyl at absorbance 420 nm from the substrate *p*-nitrophenyl-beta-D-glucoside (pNPG) at a range of substrate concentrations. The data was then fitted using the Michaelis-Menten equation to determine *k*_cat_ and K_M_.

The thermal stability (T_m_) for the mutants were determined using the Protein Thermal shift (PTS)™ kit made by Applied BioSystem ® from Thermo Fisher. Following the standard protocol by the manufacturer, purified proteins were diluted from 0.1 to 0.5 mg/mL and fluorescence reading was monitored using QuantaStudio™ 3 System from 20 °C to 90 °C. The T_M_ values were then determined using the two-state Boltzmann model from the Protein Thermal Shift™ Software 1.3 by Applied BioSystem ® from Thermo Fisher.

## RESULTS

### Molecular modeling of three mutants using FoldIt

All mutants were hypothesized to maintain similar kinetic activity based on the residues being on the surface of the enzyme and are 19-20 Å from the active-site (Fig 1). Furthermore, based on the change of total system energy (ΔTSE) between the WT and the mutant, M319C (ΔTSE of −2.55) and K337D (ΔTSE of −2.368), we predicted a slight increase in thermal stability (T_M_) compared to WT (Fig 2). On the other hand, the change from hydroxyl phenol group with hydrogen bonding capability to an alkyl group for mutant T431I we predicted a small decrease in stability (ΔTSE of +0.86).

**Figure 2.**
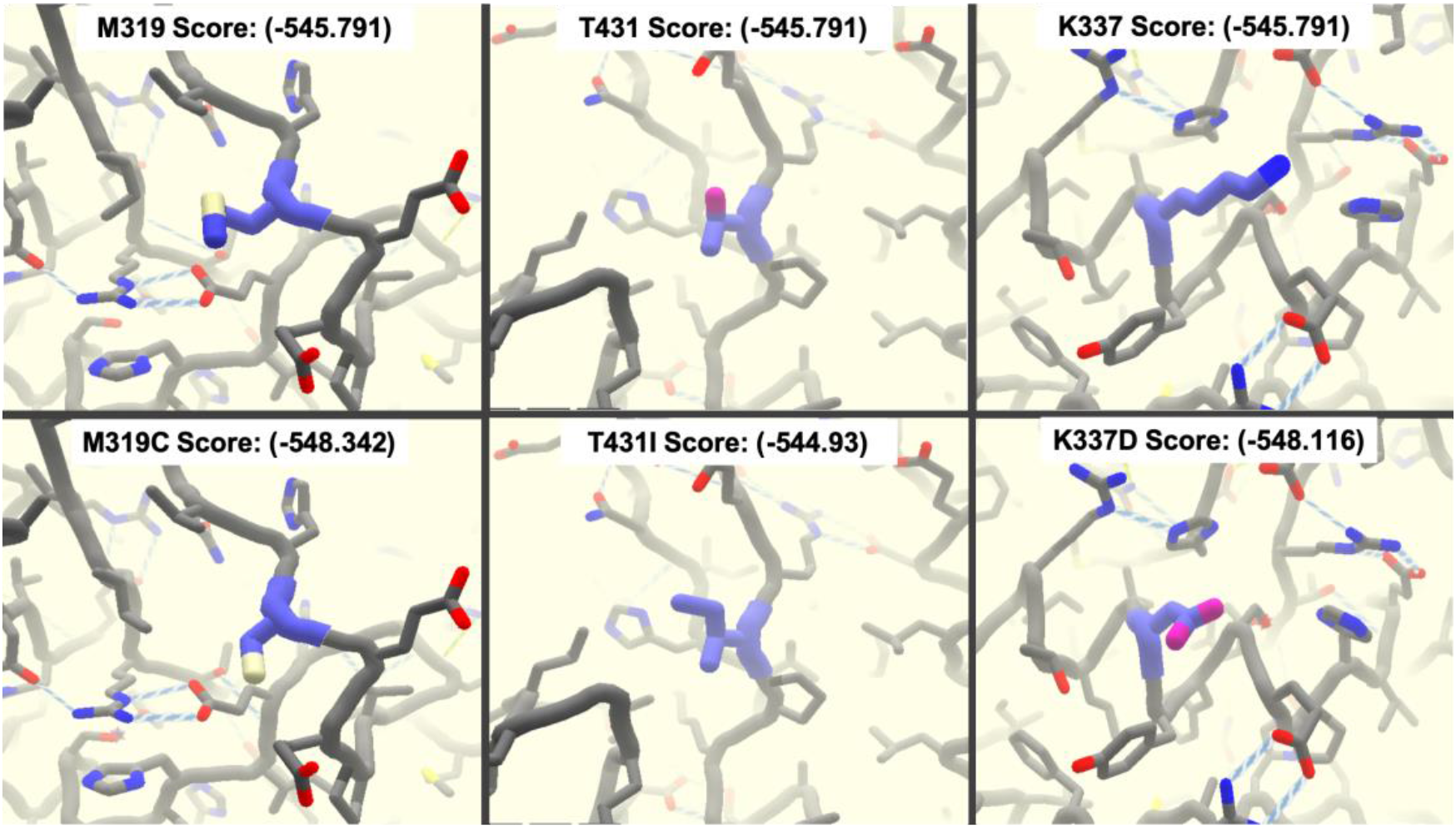
Molecular modeling of the three variants. Using the FoldIt modeling tool, the three mutants: M319C, T431I, and K337D are shown from left to right, respectively. Top row shows each mutational sites of the wild type with its initial total system energy score (TSE of −545.791). The bottom row shows the new molecular environment and TSE upon mutation.

### Protein expression and purity

All protein samples were able to express as soluble proteins as shown in Fig 3. Using 4-12% SDS-PAGE gel, the purity of all samples was greater than 80% for kinetic characterization and thermal stability assay.

**Figure 3.**
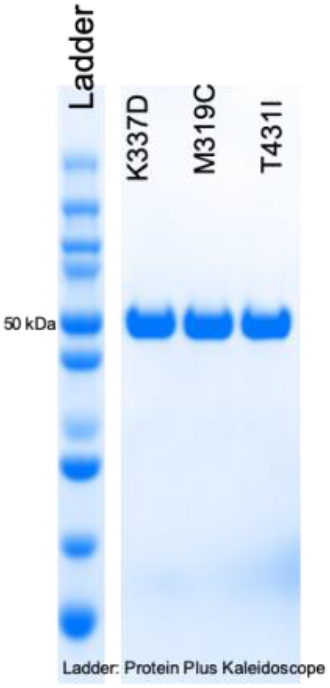
Purity and expression of three variants. The purity of all protein samples were evaluated SDS-PAGE. All samples are showing at 50 kDa from left to right: K337D, M319C, and T431I.

### Michaelis-Menten kinetic parameters

The Michaelis-Menten parameters (*k*_cat_, K_M_, and *k*_cat_/K_M_) for the three mutants were measured using the reporter substrate *p*-nitrophenyl-beta-D-glucoside (pNPG). The colorimetric assay for each mutant was done in triplicates. In this study, the *k*_cat_, K_M_, and *k*_cat_/K_M_ for wild type are 1070 ± 10 Min-1, 6.5 ± 0.2 mM, and 165 ± 35 mM-1 Min-1, respectively. All mutants exhibit no significant changes in catalytic efficiency compared to WT (SI Table 1). No significant change in K_M_ was observed for the mutants with 4.3mM, 5.1mM, and 6mM for K337D, T43ID, and M319C respectively (Fig 4). The kinetic constants between wild type and all three mutant proteins are consistent with modeling predictions of being distal from the active site and relatively small changes in the TSE.

**Figure 4.**
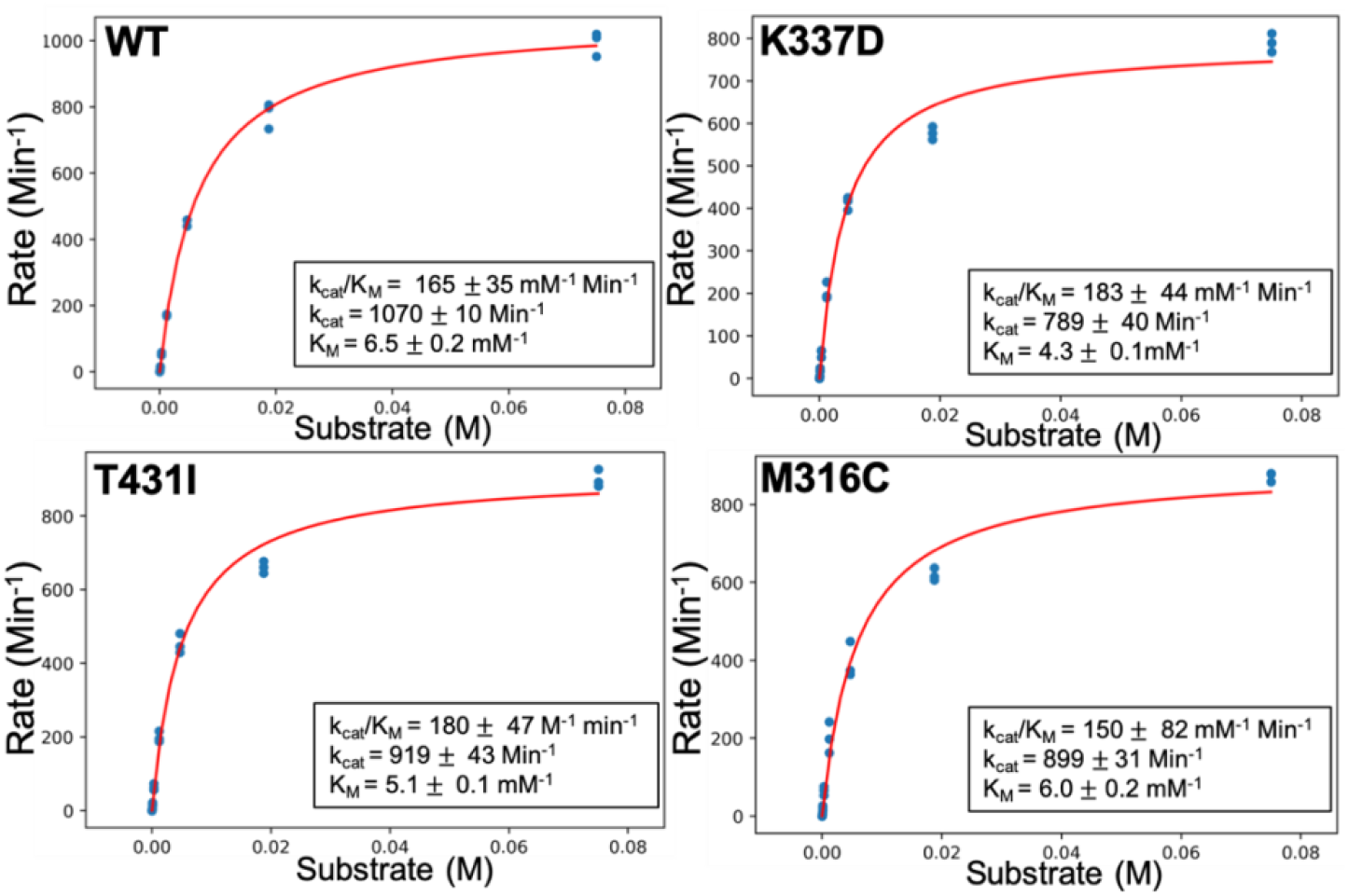
Michaelis-Menten constants (*k*_cat_, K_M_, and *k*_cat_/K_M_) of the WT and three mutants. All distal mutants showed no significant changes in kinetic activity compared to wild type which is consistent with modeling predictions.

### Thermal stability of the designed mutants

The observed T_M_ for both M319C (45.8 ± 0.2 °C) and T431I (43.6 ± 0.1) °C remained fairly close to the WT (45.3 ± 0.4 °C), both having less than a 2 °C change (Fig 5). The mutant with the most drastic change in thermal stability is K337D (38.8 ± 0.8 °C) with roughly a 6 °C decrease from WT. Based on the Foldit simulations both M319C and K337D mutants were hypothesized to be slightly more stable than WT, while T431I was predicted to be less stable due loss of hydrogen bonding interactions. These predictions are consistent with the experimental observations that M319C will have a slight increase in stability, while T431I is slightly less stable than WT. However, for mutant K337D, the prediction did not match with experimental T_M_ result since the mutant was hypothesized to have a small increase in stability.

**Figure 5.**
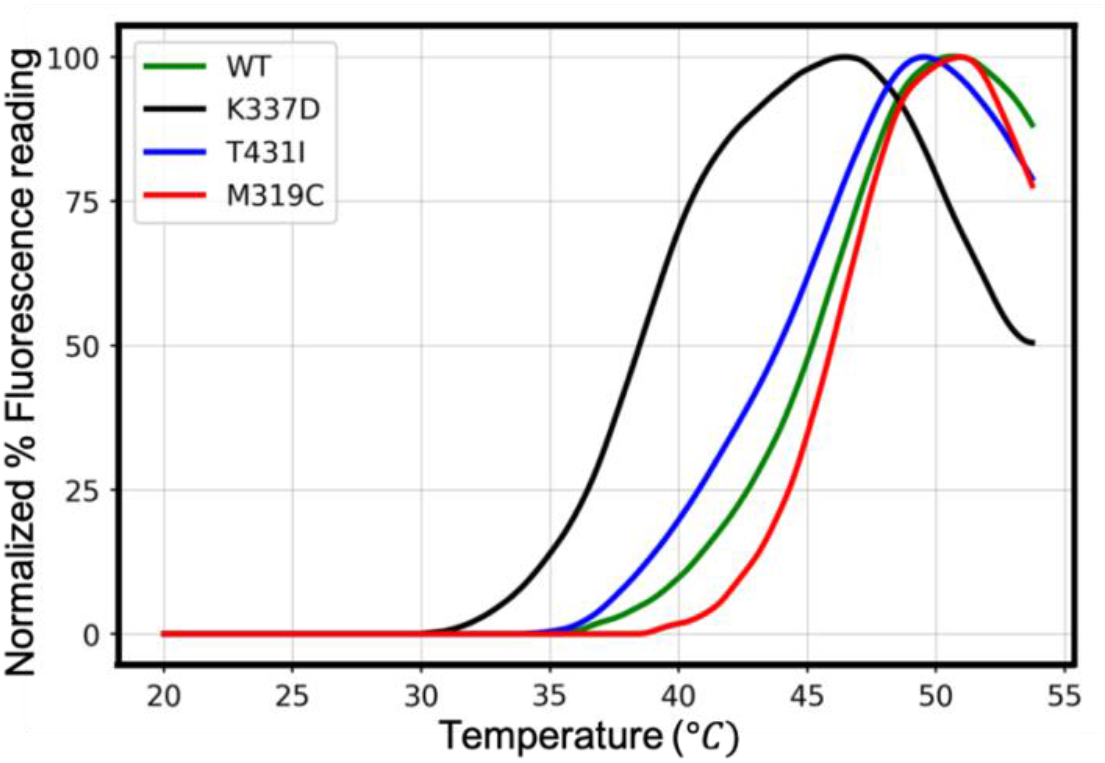
Temperature melting curves for WT and three mutants. The normalized percent fluorescence readings of the melting curve using the dye-based Protein Thermal Shift assay. The green, black, blue, and red depicts the average data of three replicates observed for WT, K337D, T431I, and M319C, respectively.

## CONCLUSION

The three mutants in this study were all chosen on the surface of the protein and distal from the active-site. These variants did not exhibit unexpected kinetic activity changes compared to WT, which supports our hypothesis due to the mutants being surface residues. Furthermore, the stability changes predicted for two mutants from FoldIt-based molecular modeling was also in agreement with measured changes in thermal stability, albeit relatively small changes. Unexpectedly, the FoldIt predictions did not match with the thermal stability results for K337D. A potential reason for the inaccurate predications could be due to the mutational changes causing an electrostatic repulsion with another nearby aspartic acid residue. The electrostatic term within the Rosetta energy function is a statically-derived approximation14 and therefore for the specific environment of K337D effect of this interaction may have been underestimated.

Overall, this data supports efforts to focus on mutations in either the core or interface of the enzyme-substrate complex to obtain the most significant changes. However, for subtle improvements, optimization of surface residues should not be overlooked as the additive effects of M319C could potentially increase thermal stability by almost 1°C. More importantly, these mutants continue to add to the Design to Data efforts to quantitatively catalog the effects of a mutation on BglB in a consistent and systematic fashion. Datasets such as these have the potential to be the foundation for developing new algorithms that accurately predict a mutations’ effect on protein thermal stability and kinetic activity.

---

**SI Table 1.** Total system energy (TSE), kinetic parameters, and thermal stability (T_M_) of WT and three distal mutants.

| Protein | TSE | $k_{cat}$<br>(Min<sup>-1</sup>) | $K_M$<br>(mM) | $k_{cat}/K_M$<br>(mM<sup>-1</sup> Min<sup>-1</sup>) | $T_M$ (°C) |
| --- | --- | --- | --- | --- | --- |
| WT | -545.79 | 1070 ± 10 | 6.5 ± 0.2 | 165 ± 35 | 45.3 ± 0.4 |
| M319C | -548.34 | 899 ± 31 | 6.0 ± 0.5 | 150 ± 82 | 45.8 ± 0.2 |
| T431I | -544.93 | 919 ± 43 | 5.1 ± 0.1 | 180 ± 47 | 43.6 ± 0.1 |
| K337D | -545.12 | 789 ± 40 | 4.3 ± 0.1 | 183 ± 44 | 38.8 ± 0.8 |

The estimated errors are the s.d of three independent measurements.

